# Can recent evolutionary history promote resilience to environmental change?

**DOI:** 10.1101/2022.11.04.515151

**Authors:** Eleanor Kate Bladon, Sonia Pascoal, Rebecca Mary Kilner

## Abstract

Principles of social evolution have long been used retrospectively to interpret social interactions, but have less commonly been applied predictively to inform conservation and animal husbandry strategies. We investigate whether differences in developmental environment, facilitated by divergent social conditions, can predict resilience to environmental change. Upon exposure to harsh novel environments, populations that previously experienced more benign social environments are predicted either to suffer fitness losses (the “mutation load hypothesis” and “selection filter hypothesis”) or maintain fitness (the “beneficial mutation hypothesis”). We tested these contrasting predictions using populations of burying beetles *Nicrophorus vespilloides* we had evolved experimentally for 45 generations under contrasting social environments by manipulating the supply of post-hatching parental care. We exposed sexually immature adults from each population to varying heat stress and measured the effect on survival and reproduction. The greater the level of parental care previously experienced by a population, the better its survival under heat stress during sexual maturation. Although this is consistent with the “beneficial mutation hypothesis”, it is also possible that populations that had evolved without post-hatching care were simply more prone to dying during maturation, regardless of their thermal environment. Overall, we suggest that stochastic genetic variation, probably due to founder effects, had a stronger influence on resilience. We discuss the implications for translocation and captive breeding programmes.

**Lay summary:** Can we use knowledge of a population’s evolutionary history to predict how individuals might cope with environmental change? We investigated whether burying beetles that had evolved for 45 generations with or without post-hatching parental care differed in their resilience to extreme temperatures as they developed to sexual maturity. We found limited evidence that experimental evolution under different regimes of parental care contributed to thermal resilience, which instead was better explained by chance historical events.

## Introduction

In a world changing rapidly through the effects of climate change (Pörtner et al., 2022), habitat degradation (Millenium Ecosystem Assessment, 2005) and excessive use of antibiotics (Graham et al., 2019) and pesticides (Hawkins et al., 2019), our ability to predict how effectively organisms will adapt to new environments is more important than ever. The principles of social evolution, and how they influence developmental environment, have long been used retrospectively to understand social interactions (Bourke, 2011). However they have less commonly been used to inform the development of practical strategies to address global biological challenges (Carroll et al., 2014). Yet an understanding of a population’s social evolutionary history, by which we mean the type of selective social environment it has experienced, could predict its resilience to future environmental perturbation, through the effects that different social environments have on mutation load and selection for particular alleles (Lumley et al., 2015). This has practical value in its potential to address several major economic and societal problems, including food security (pest/crop interactions, crop development), human and animal medicine (antibiotic development, anti-parasite interventions) and wild animal conservation (choice of populations for translocation schemes, identification of populations vulnerable to disease and environmental change) (Carroll et al., 2014).

In theory, historical effects on resilience to the current environment depend on whether the past developmental environment experienced by the population is benign or harsh - because this influences the extent of standing genetic variation available. Few studies have tested experimentally how past exposure to different social environments during development affects a population’s future response to environmental change (but see Lumley et al., 2015).

Benign social interactions, which are usually due to some form of cooperative interaction, are known to buffer individuals from environmental stress and therefore to reduce the extent to which mildly deleterious alleles are exposed to natural selection (Linksvayer & Wade, 2009; Mashoodh et al., 2023; Pascoal et al., 2022). Elevated mutation loads resulting from relaxed selection have been shown to reduce reproductive success in animals kept in captive breeding programmes (Araki et al., 2007; McPhee, 2004; Robert, 2009) and have been linked to a higher prevalence of genetic disorders in human societies where selection is buffered by widely available healthcare (Lynch, 2016; Saniotis et al., 2021; You & Henneberg, 2018). The “mutation load hypothesis” therefore predicts that individuals previously exposed to more benign developmental environments, facilitated by the buffering effect of social environments which impose fewer selective pressures, should be less resilient to new environmental stresses. However, whilst most of the mutations that accumulate under relaxed selection are likely to be (mildly) harmful, some could potentially be beneficial and might even facilitate greater resilience to environmental change at the population level. The “beneficial mutation hypothesis” therefore conversely predicts that individuals that develop in more benign social environments could be more resilient to new environmental stresses, if by chance they have accumulated cryptic genetic variation that is beneficial when expressed in these new environments (Barrett & Schluter, 2008; Paaby & Rockman, 2014). At the other end of the spectrum, a harsh social environment potentially imposes a strong selection filter, which is likely to reduce the population’s mutation load, causing individuals generally to be fitter (Balmford, 1996). The “selection filter hypothesis” predicts that a history of exposure to harsher social environments means that populations are more likely to be resilient to environmental change.

Species in which offspring are facultatively dependent on parental care (Clutton-Brock, 1991) offer a way to test these contrasting predictions. Offspring that receive care are exposed to a benign social environment during development which, if sustained over many generations relaxes selection (Snell-Rood et al., 2016). Conversely, offspring that receive no care over many generations experience a harsher environment during development and consequently experience stronger selection against deleterious mutations. Our experiments focus on the burying beetle *Nicrophorus vespilloides*, which exhibits elaborate biparental care upon which offspring are facultatively dependent (Capodeanu-Nägler et al., 2016). We know from previous work that offspring exposed to post-hatching care experience a more benign developmental environment than those that develop without post-hatching care. For example, Schrader et al. (2017) found that when breeding pairs of beetles were removed before providing post-hatching care, fewer of them produced at least one surviving larva, and those larvae that did survive were lighter. We exploited this natural history to test whether, and how, past social interactions influence future resilience to environmental change.

Burying beetle adults locate the carcasses of small vertebrates using their highly sensitive antennae (Kalinová et al., 2009; Trumbo & Steiger, 2020) and then fight intrasexually for ownership (Otronen, 1988). The winning pair then tears the fur or feathers from the carcass, covers the flesh with antimicrobial oral and anal exudates, rolls it into a ball and buries it underground (Cotter & Kilner, 2010). The female then lays eggs in the soil surrounding the carcass nest. When the larvae hatch, 2-3 days later, they crawl to the nest (Müller & Eggert, 1989), where their parents defend them from predators and intra- and interspecific competitors and feed them trophallactically (Eggert et al., 1998). The duration of post-hatching care is highly variable and in some cases is non-existent due to the death or desertion by parents after preparation of the carcass nest (Schrader, Jarrett, et al., 2015a; Smiseth et al., 2003). In such cases larvae can self-feed (Smiseth et al., 2003) and can survive (at least in the laboratory) without any post-hatching care.

We evolved two replicate populations of burying beetles in the laboratory under contrasting regimes of parental care for 45 generations. In the Full Care populations, beetles were allowed to stay with their offspring throughout development. Individuals in these populations experienced a more benign environment during development and relaxed selection (Pascoal et al., 2022). By contrast, in the No Care populations, parents were removed just prior to larval hatching and so were unable to supply any post-hatching care at all. Therefore individuals in these populations experienced a harsher social environment and stronger selection against deleterious mutations (Pascoal et al., 2022).

After 45 generations of experimental evolution, we tested whether there were differences in the resilience of beetles from these two types of experimental population when they were exposed to higher temperatures during their juvenile adult stage. We define resilience as the ability of a population to maintain normal processes, in particular reproductive success, in response to a harsh novel environment (Lowerre-Barbieri et al., 2017; Martin & Wiebe, 2004; Wingfield et al., 2011). Temperature was chosen as the environmental stressor because of its ecological relevance in the context of climate change. Insects, being poikilothermic, are particularly vulnerable to heat stress, and research on burying beetles specifically has shown that they have reduced breeding success at higher temperatures. We chose to expose the beetles to the temperature treatments during sexual maturation alone because this is the time when an adult beetle’s gonads are developing (e.g. David et al., 2005; Gibbs et al., 2010; Sales et al., 2018; Zizzari and Ellers, 2011). Furthermore, Sales et al. (2021) show in flour beetles *Tribolium castaneum* that this developmental stage is particularly vulnerable to effects of thermal stress on offspring production. Additionally, the natural history of the burying beetle will likely render this a particularly sensitive period as sexually immature individuals cannot stay underground to buffer themselves from heat stress because they must emerge to feed.

We exposed sexually immature adults to different degrees of heat stress and measured the effect on the following correlates of fitness: survival to sexual maturity, anal exudate lytic activity, carcass sphericity, fecundity, and hatching success Duarte et al., 2021). We predicted that the contrasting regimes of selection imposed by these different social environments during development for the 44 preceding generations should yield a contrasting response to heat stress during sexual maturation. The “mutation load hypothesis” and “selection filter hypothesis” each predict that individuals derived from the No Care populations should be more resilient to heat stress, whereas the “beneficial mutation hypothesis” predicts that individuals derived from the No Care populations should be less resilient to heat stress.

## Methods

### Burying beetle husbandry in the laboratory: general methods

Burying beetles were bred by placing them in a plastic breeding box (17 x 12 x 6 cm), partially filled with damp soil (Miracle-Gro Compost), furnished with a 10-15 g mouse carcass for breeding. Boxes were placed in a cupboard after pairing to simulate underground conditions. Larvae were counted and weighed 8 days after pairing and each one placed in its own cell (2 x 2 x 2 cm) within a plastic pupation box (10 x 10 x 2 cm), filled with damp peat. Sexually immature adults were eclosed around 21 days later and each was placed in its own individual box (12 x 8 x 2 cm). Adults were fed twice weekly with beef mince until breeding, which took place 15 days post-eclosion. At all life stages, individuals were kept at 21 °C. During breeding, pairs were kept in a constantly dark environment, but for all other life stages they were kept on a 16L: 8D hour light cycle.

### Experimental Evolution

The *N. vespilloides* populations used in this study were part of a long-term experimental evolution project that investigated how populations of burying beetles adapt to the loss of parental care. There were four experimental populations in all: Full Care (FC) (replicated twice) and No Care (NC) (replicated twice). These populations were established from a stock population founded with wild-caught beetles trapped under permit in 2014 from four woodland sites across Cambridgeshire, UK (Byron’s Pool, Gamlingay Woods, Waresley Woods and Overhall Grove). From this stock population 32 pairs were bred, with the male and female coming from different sites to minimise the likelihood of inbreeding. The resulting 671 offspring established the experimental evolution populations. FC populations were left undisturbed (as described in general methods above) and allowed to stay with their offspring until dispersal. However, in the NC populations, at 53 h post-pairing, when the carrion nest was complete but before the larvae had hatched, both parents were removed to prevent post-hatching care. This procedure was repeated at every generation. The replicate experimental populations were bred in two separate blocks (FC1/NC1 and FC2/NC2), and the timing of the lifecycle was staggered between blocks to be 7 days apart. As all the blocks were derived from the same stock population, FC1 and NC1 were no more related to each other than FC2 and NC2.

### Testing for resilience to heat stress

#### Experimental generation 1: Partitioning current care environment from evolutionary history of care

Beetles were taken for analysis at generation 45 of experimental evolution. We partitioned the effects of current care experienced by individuals (No Care or Full Care), from an evolutionary history of care experienced by the source population (No Care or Full Care), by breeding beetles for one generation using the following treatments:

1) Adults from Full Care populations allowed to provide full parental care (Full Care parents with a Full Care regime) = 73 FC_FC_ pairs
2) Adults from Full Care populations removed at 53 hours after pairing (Full Care parents with a No Care regime) = 74 FC_NC_ pairs
3) Adults from No Care populations removed at 53 hours after pairing (No Care parents with a No Care regime) = 74 NC_NC_ pairs
4) Adults from No Care populations allowed to provide full parental care (No Care parents with a Full Care regime) = 74 NC_FC_ pairs

When larval development was complete, and larvae were starting to disperse away from the remains of the carrion nest (8 days after pairing), larvae were put in pupation boxes to mature and reach eclosion, as described above.

#### Experimental generation 2: Exposure to heat stress

Immediately after eclosion, each experimental individual was placed alone in a small plastic box (12 x 8 x 6 cm) containing a fine layer of soil. The boxes were placed in incubators (Panasonic MLR-352H-PE) set to either 22 °C, 25 °C or 28 °C on a 16L: 8D hour light cycle. The maximum temperature was chosen based on a previous, unpublished pilot experiment in the laboratory that found a severe reduction in hatching success of beetles reared at 28 °C, and the 25 °C temperature was chosen to represent an intermediate temperature between one that was likely to cause severe heat stress and one that was close to our standard laboratory temperature. Our intention was not to mimic natural thermal conditions closely, but rather create different levels of heat stress across the treatments within a naturally occurring range. The boxes were removed from the incubators briefly once after a week so that the beetles could be fed with beef mince. We then measured the following correlates of fitness for this generation of beetles:

#### Survival

To measure the effect of heat stress on survival, after 14 days (at sexual maturity) all individuals were removed from their incubators and classified as alive or dead.

The surviving individuals were then assigned to one of the following treatments for breeding:

1) 22 °C male x 22 °C female (n = 29 FC_FC_, 29 FC_NC_, 28 NC_NC_, 29 NC_FC_)
2) 25 °C male x 25 °C female (n = 36 FC_FC_, 35 FC_NC_, 36 NC_NC_, 36 NC_FC_)
3) 28 °C male x 28 °C female (n = 32 FC_FC_, 34 FC_NC_, 29 NC_NC_, 30 NC_FC_)

Individuals were only bred within their populations and temperatures, e.g. 22 °C males from FC_FC_ were only bred with 22 °C females from FC_FC_, resulting in 12 treatments (Figure S1).

Pairs were placed in breeding boxes containing 350 ml of moist soil and a 10-13 g mouse carcass, and the box was sealed and kept in a dark environment, as described under ‘general methods’ above. All pairs were kept at the standard laboratory temperature of 21 °C, regardless of their temperature treatment, to isolate the effects of the thermal environment during sexual maturation on subsequent reproductive success.

#### Lytic activity

Higher exudate lytic activity is heritable and increases larval survival so it is possible that there has been differential selection for this during the evolution of the experimental lines (Arce et al., 2012; Cotter & Kilner, 2010). Additionally, previous research has shown that when mature adults of other *Nicrophorus* species were subjected to a three-day thermal stress treatment there were significant contrasting effects on the lytic activity of their oral exudates (Jacques et al., 2009). Therefore, we would expect to see lower levels of lytic activity in beetles more negatively affected by higher temperatures.

At 53 h after pairing, we collected anal exudates from parents from a subsample of broods from each treatment (n = 27 FC_FC_, 27 FC_NC_, 22 NC_NC_, 19 NC_FC_) by holding each beetle by the abdomen and pressing a 5 ul glass microcapillary tube (Drummond) to its anus. Each sample was pipetted immediately into a 1.5 ml Eppendorf tube on dry ice and subsequently stored at −80 °C until analysis. We quantified the lytic activity of each sample in an automated microplate reader (BioTek ELx808) by running turbidity assays that measured the degradation of bacterial cell walls (see Duarte et al., 2021 for full details). Briefly, 1ul of each exudate sample was diluted in 24 ul of 6.4 pH 0.02 M potassium phosphate buffer and 10ul of this solution was added to a well of a 96-well flat-bottomed microplate (Thermo Scientific Nunc MicroWell) containing 100 ul of a 1.3 mg ml^-1^ solution of lyophilized *Micrococcus lysodeikticus* cells in potassium phosphate buffer. The microplate reader was set to 25 °C and took an initial reading at 450 nm. Plates then went through a cycle of being shaken at a medium setting every 10 minutes, followed immediately by a 450 nm reading, for 60 minutes.

#### Carcass sphericity

Rounder carcasses are less hospitable to parasitism by blowflies (Sun & Kilner, 2020), and are negatively correlated with parental lifespan, indicating that preparing a rounded carcass requires significant resources from the parents (De Gasperin et al., 2016). Therefore, we would expect that if higher temperatures have a negative effect on beetles then they would make a less spherical carcass.

At 53 h, we also briefly removed the carcass from a subsample of broods (n = 33 FC_FC_, 31 FC_NC_, 33 NC_NC_, 33 NC_FC_) where the parents had experienced the 22 °C and 28 °C degree treatments. Each carcass was placed on a white background, alongside a scale for measurement, and photographed from perpendicular angles using two cameras (Canon DSLR, with 18-55mm lens) at 30 cm away from the carcass (Figure S2).

#### Laying failure, clutch size, hatching failure and brood size

At 53 h after pairing, we removed the parents from all 383 broods and counted the number of eggs laid, through inspection of the underside of the box (following Jarrett et al., 2017; Schrader et al., 2015c). At 70, 80, 94 and 104 h after pairing we checked the carcass and the surface of the soil for any first instar larvae, and counted them before discarding them.

### Statistical analyses

All statistical tests were conducted in R version 3.6.1 (R Core Team, 2022) and a 0.05 significance threshold was used throughout. Data handling and visualisation were carried out using base R and the ‘tidyverse’ suite of R packages (Wickham et al., 2019). Model selection for the analyses of brood mass and brood size was conducted using the ‘dredge’ and ‘model.avg’ functions in the ‘MuMIn’ R package (Bartoń, 2023). The most parsimonious model within two AIC_C_ points of the model with the lowest AIC_C_ was chosen as the optimal model. Goodness of fit of optimal models was assessed with the ‘DHARMa’ package (Hartig, 2022) or, in the case of beta regression models, the base ‘stats’ package. Continuous independent variables in all models were scaled and centred using the ‘scale’ function in base R.

#### Survival

We ran a binomial GLM with survival status as the dependent variable (“0” = died in the incubator, “1” = survived to sexual maturity), and population evolutionary history (Full Care or No Care), generation 1 care regime (Full Care or No Care), temperature, sex, block, and the two-way interactions of these terms, and three-way interactions between population evolutionary history, block and temperature, and generation 1 care regime and history, block and temperature as the independent variables in the maximal model. See Table S1 for an explanation of why each interaction term was included in the models.

#### Lytic activity

In some of the broods only one of the parents produced enough exudate to allow us to conduct the lytic activity analysis on their sample. The total useable sample size for exudates was 150 individuals, and for 110 of these we had a sufficient sample from their partner (55 pairs) and in the other 40 only one partner in the pair could be analysed (we could not collect enough exudate from the partners of 20 females and 20 males used in the analysis).

We used the percentage change between the 450 nm absorption at the first reading (0 minutes) and at 60 minutes as the dependent variable (as per Duarte et al., 2021). We first ran a paired t-test to determine whether there was a consistent sex difference within pairs in lytic activity. As the lytic activity of one individual in a pair may influence its partner’s levels, for males and females we then ran separate beta regression, using the ‘betareg’ R package (Cribari-Neto & Zeileis, 2010), with proportion of cells degraded (450 nm reading at 60 minutes/450 nm reading at 0 minutes) in either males’ or females’ anal exudate as the dependent variable, and population evolutionary history (Full Care or No Care), generation 1 care regime (Full Care or No Care), temperature, partner’s lytic activity, block, the two-way interactions between temperature and evolutionary history, and sex and temperature, and the three-way interaction between population evolutionary history, block and temperature, and generation 1 care regime, block and temperature as independent variables in the maximal model.

#### Carcass sphericity

Carcass sphericity was calculated digitally in ImageJ (Schneider et al., 2012), using a macro script adapted from De Gasperin et al (2016) (Script S1 and Figure S3). The ImageJ script assumes that if a carcass were perfectly spherical, the two-dimensional images in the carcass from the top and side would be exact circles. The script compares the actual images to a perfect circle of the same perimeter and then calculates the difference between the two – such that a perfectly spherical carcass would have a score of 1 to denote that it fills 100% of the space of a perfect sphere (Figure S4).

As carcass sphericity was bounded between 0 and 1, we ran a beta regression with carcass sphericity percentage as the dependent variable and population evolutionary history (Full Care or No Care), generation 1 care regime (Full Care or No Care), temperature, sex, block, and the two-way interactions of these terms, and three-way interactions between population evolutionary history, block and temperature, and generation 1 care regime and history, block and temperature as the independent variables in the maximal model.

#### Laying failure, clutch size, hatching failure and brood size

To investigate predictors of fecundity, we first ran a binomial GLM with laying failure (“0” = for a brood where no eggs were laid and “1” = for a brood where at least one egg was laid) as the dependent variable and population evolutionary history (Full Care or No Care), generation 1 care regime (Full Care or No Care), temperature, carcass mass, block, two-way interactions between population evolutionary history and generation 1 care, population evolutionary history and temperature, generation 1 care and temperature, population evolutionary history and block, generation 1 care and block, and block and temperature, and three-way interactions between population evolutionary history, block and temperature, and generation 1 care, block and temperature as independent variables.

Then, to examine predictors of clutch size in the broods that did lay eggs, we ran a quasi-Poisson GLM with clutch size (the number of eggs counted from the underside of the breeding box) as the dependent variable, and the same independent variables that were used in the model analysing laying failure vs success.

We then ran two tests to investigate predictors of offspring production. Firstly, we ran a binomial GLM with brood success (“0” = for a brood with no larvae, “1” = for a brood with at least one hatching larva) as the dependent variable and the same independent variables that were used in the binomial model analysing laying failure vs success. Next, for broods where at least one egg hatched, we ran a quasi-Poisson GLM with brood size (number of larvae counted on the carcass or soil) as the dependent variable and the same independent variables that were used in the model analysing laying failure vs success, plus clutch size. Broods were included in the hatching success model even if no eggs were counted from the underside of the breeding box, in case by chance a female had laid very few eggs more shallowly in the soil which then successfully hatched.

## Results

### Survival

The optimal model retained population evolutionary history, temperature, block (evolutionary population replicate) and the interaction between temperature and block (Table 1 and S2). Beetles from populations with an evolutionary history of No Care were more likely to die in the incubators, regardless of the temperature they experienced in the incubator or the care they received in generation 1 of the heat stress experiment. Mean survival of Full Care individuals across blocks was 94%, but 87% for No Care individuals.

**Table 1.** Results of binomial model investigating the effects on likelihood of a beetle surviving from eclosion to sexual maturity in incubators set to 22 °C, 25 °C or 28 °C. Terms retained in the optimal model based on AIC_C_ are presented.

| Independent variable | Estimate | SE | z | p |
| --- | --- | --- | --- | --- |
| Intercept | 4.170 | 0.301 | 13.857 | <0.001 |
| Population evolutionary history (No Care) | -1.008 | 0.231 | -4.362 | <0.001 |
| Temperature | -0.335 | 0.255 | -1.315 | 0.189 |
| Block (2) | -1.041 | 0.341 | -3.055 | 0.002 |
| Block:temperature | -1.490 | 0.343 | -4.339 | <0.001 |

To further investigate how block affected survival at higher temperatures we split the dataset by block and ran generalised linear models with survival probability as the dependent variable and temperature as the independent variable. In block 1 the impact of temperature was non-significant (χ^2^_1, N=618_ = 1.820, *p* = 0.177), but in block 2 there was a significant negative effect (χ^2^_1, N=600_ = 125.63, *p* = <0.001). This indicates that beetles exposed to greater heat stress as they matured sexually were more likely to die, but only in block 2.

### Lytic activity

There was no difference between males and females in lytic activity strength: t_54_ = 0.446, *p* = 0.658. However, to take partner lytic activity into account, we still ran separate models for males and females. Block was retained in the optimal models of both male and female lytic activity, with individuals having stronger lytic activity in block 2 (Figure 2, Tables 2 and S3). In the female model, temperature was also retained, with females having lower lytic activity in their exudates when they developed at higher temperatures. In the male model, generation 1 care regime was also retained, with males that received no post-hatching care having exudates with lower lytic activity. However, to test the influence of an outlier we re-ran the analysis with the removal of one datapoint where <55% of *Micrococcus* remained, and the generation 1 care term was no longer retained, indicating that the effect of generation 1 care regime was very small (Figure S5, Tables S4 & S5).

**Figure 1.**
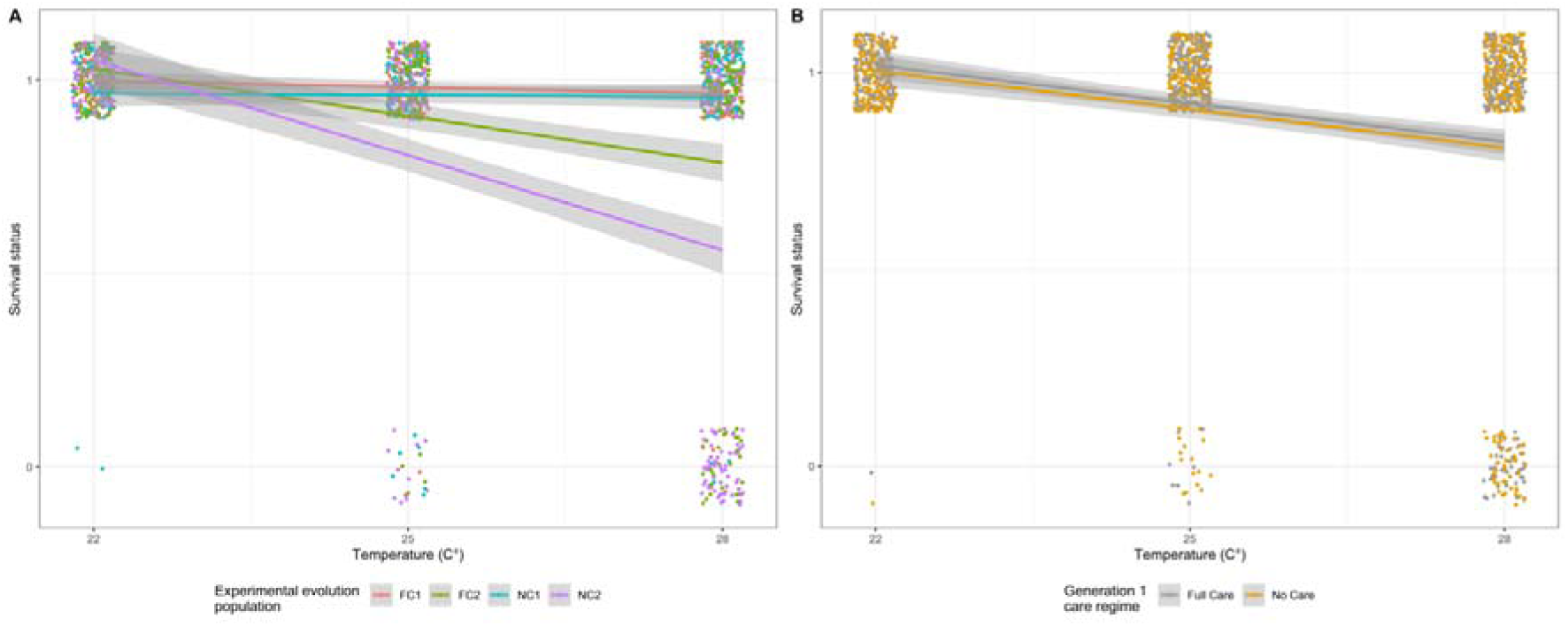
The survival of beetles kept in incubators at different levels of heat stress throughout sexual maturation (1 = survived, 0 = died) in relation to A) population evolutionary history and experimental block and B) generation 1 care regime. Points represent data for individual pairs (jittered on the x- and y-axes), lines represent the reaction norms for each treatment and the shaded areas indicate the 95 % confidence intervals. N = 1,281 individuals.

**Figure 2.**
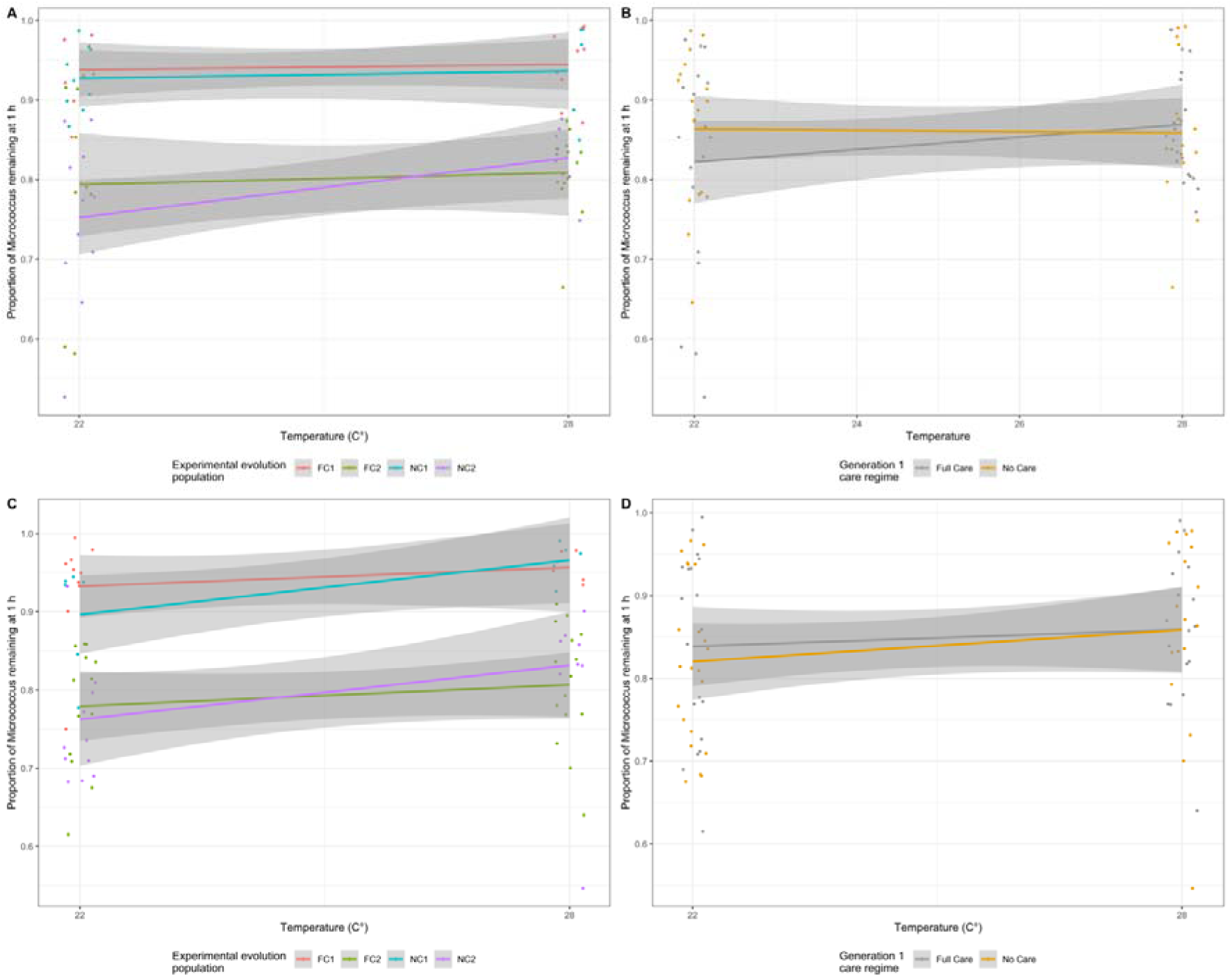
The effect of developmental temperature on male (A & B) and female (C & D) anal exudates in relation to temperature and population evolutionary history and experimental block (A & C) and temperature and generation 1 care regime (B & D). Lytic activity is defined as the percentage of cell degradation of Micrococcus lysodeikticus as recorded at 450 nm after 60 minutes. Higher remaining percentages of M. lysodeikticus remaining indicate lower exudate lytic activity. Each point represents one beetle (jittered on the x- and y-axes), lines represent the reaction norms for each treatment and the shaded areas indicate the 95 % confidence intervals. N = 75 males and 75 females.

**Table 2.**
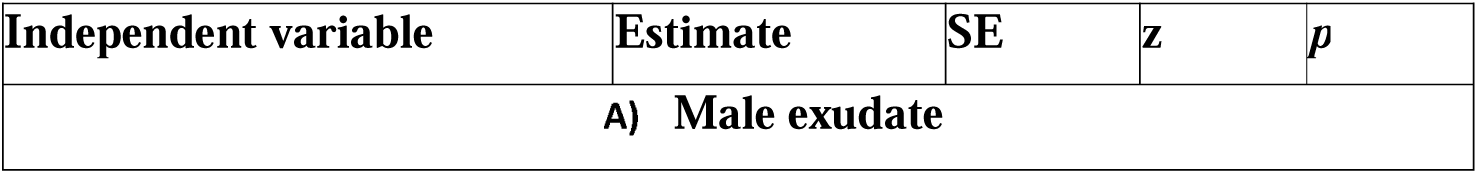
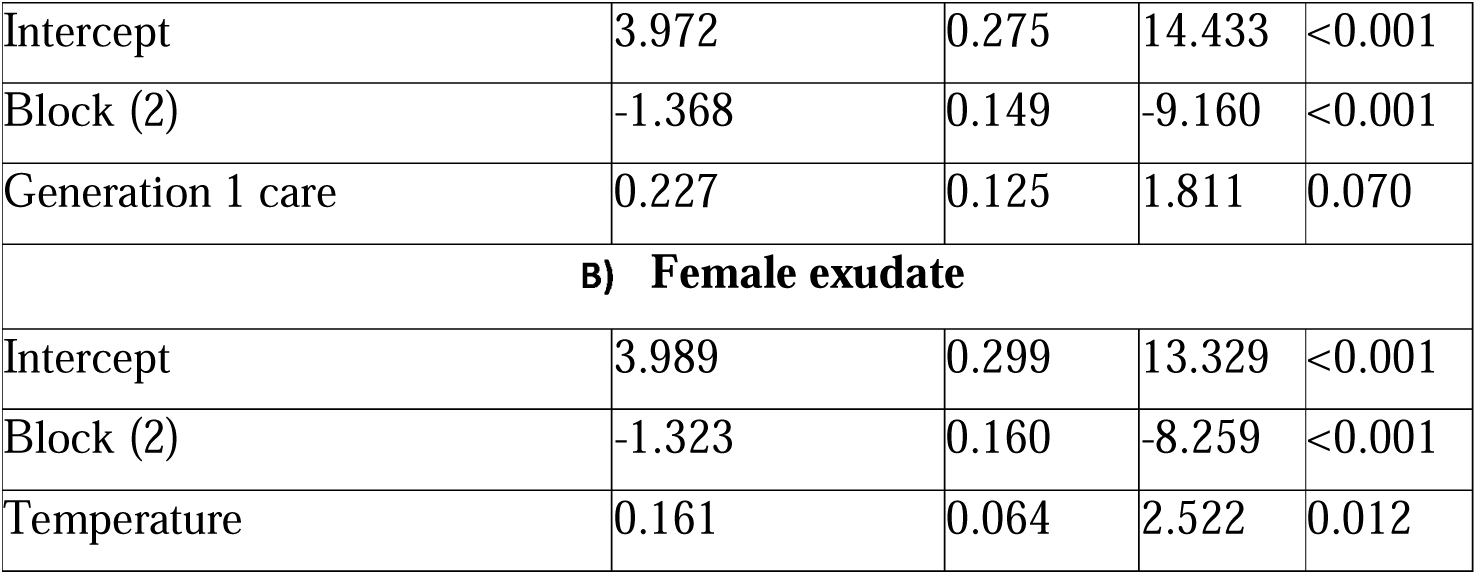
Results of beta regressions investigating the effects on A) male and B) female exudate lytic activity at 53h after pairing. Terms retained in the optimal model based on AIC_C_ are presented. We have defined lytic activity as the proportion of cell degradation of Micrococcus lysodeikticus as recorded at 450 nm after 60 minutes.

| Independent variable | Estimate | SE | z | p |
| --- | --- | --- | --- | --- |
| A) Male exudate |  |  |  |  |
| Intercept | 3.972 | 0.275 | 14.433 | <0.001 |
| Block (2) | -1.368 | 0.149 | -9.160 | <0.001 |
| Generation 1 care | 0.227 | 0.125 | 1.811 | 0.070 |
| **B) Female exudate** |  |  |  |  |
| Intercept | 3.989 | 0.299 | 13.329 | <0.001 |
| Block (2) | -1.323 | 0.160 | -8.259 | <0.001 |
| Temperature | 0.161 | 0.064 | 2.522 | 0.012 |

### Carcass sphericity

Generation 1 care regime and temperature were retained in the optimal model. Beetles that received full care prepared rounder carcasses, as did beetles that developed at 22 °C, compared to 28 °C (Figure 3, Tables 3 & S6).

**Figure 3.**
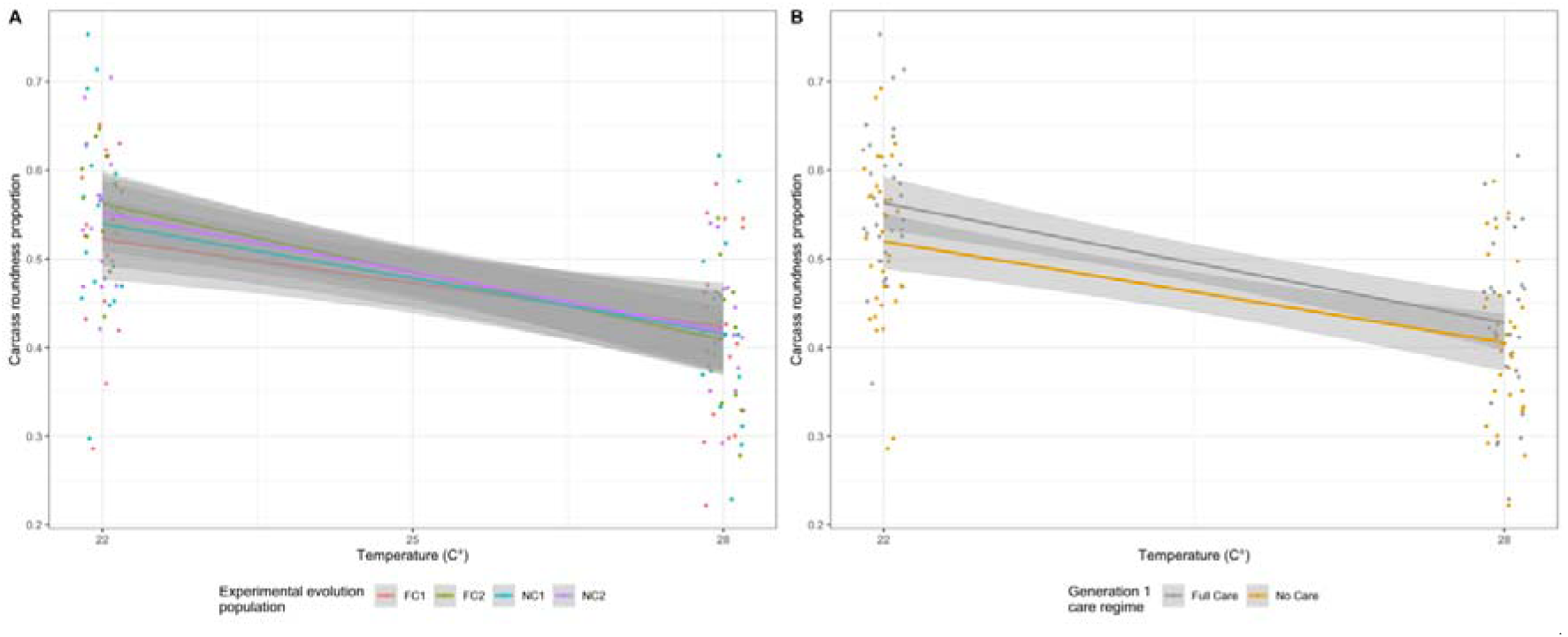
The roundness of carcasses at 53 h after pairing in relation to temperature and A) population evolutionary history and experimental block and B) generation 1 care regime. A proportion of 1 would represent a perfectly spherical carcass nest. Points represent data for individual pairs (jittered on the x- and y-axes), lines represent the reaction norms for each treatment and the shaded areas indicate the 95 % confidence intervals.

**Table 3.** Results of beta regression investigating variation in carcass nest roundness as a proxy for carcass preparation success. Terms retained in the optimal model based on AIC_C_ are presented.

| Independent variable | Estimate | SE | z | p |
| --- | --- | --- | --- | --- |
| Intercept | -0.011 | 0.044 | -0.242 | 0.809 |
| Generation 1 care | -0.134 | 0.063 | -2.145 | 0.032 |
| Temperature | -0.252 | 0.032 | -8.009 | <0.001 |

### Laying failure and clutch size

Exposure to higher temperatures during sexual maturation was more likely to cause complete failure to lay, regardless of evolutionary history, and there was a greater chance of complete laying failure in block 2 (Tables 4A & S7, Figure 4). Neither the population’s evolutionary history of care (Full Care or No Care), nor the care regime experienced in the first generation of the heat stress experiment, were retained in the optimal model of clutch success vs failure or clutch size (Tables 4B & S8). In broods where some laying occurred, higher temperatures led to fewer eggs laid per clutch, particularly in Block 1 (Figure 5).

**Figure 4.**
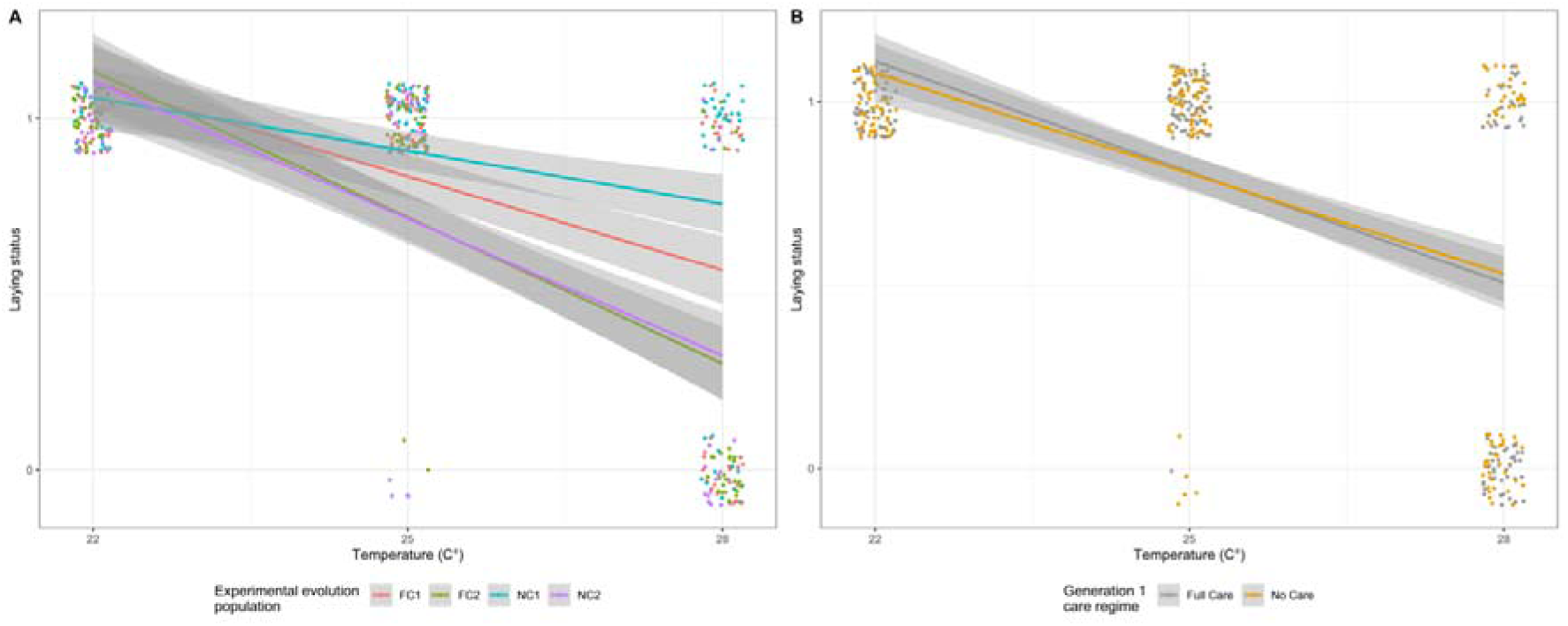
The laying status (1 = at least 1 egg laid, 0 = no eggs laid) of pairs of beetles kept in incubators at different levels of heat stress throughout sexual maturation in relation to A) population evolutionary history and experimental block and B) generation 1 care regime. Points represent data for individual pairs (jittered on the x- and y-axes), lines represent the reaction norms for each treatment and the shaded areas indicate the 95 % confidence intervals.

**Figure 5.**
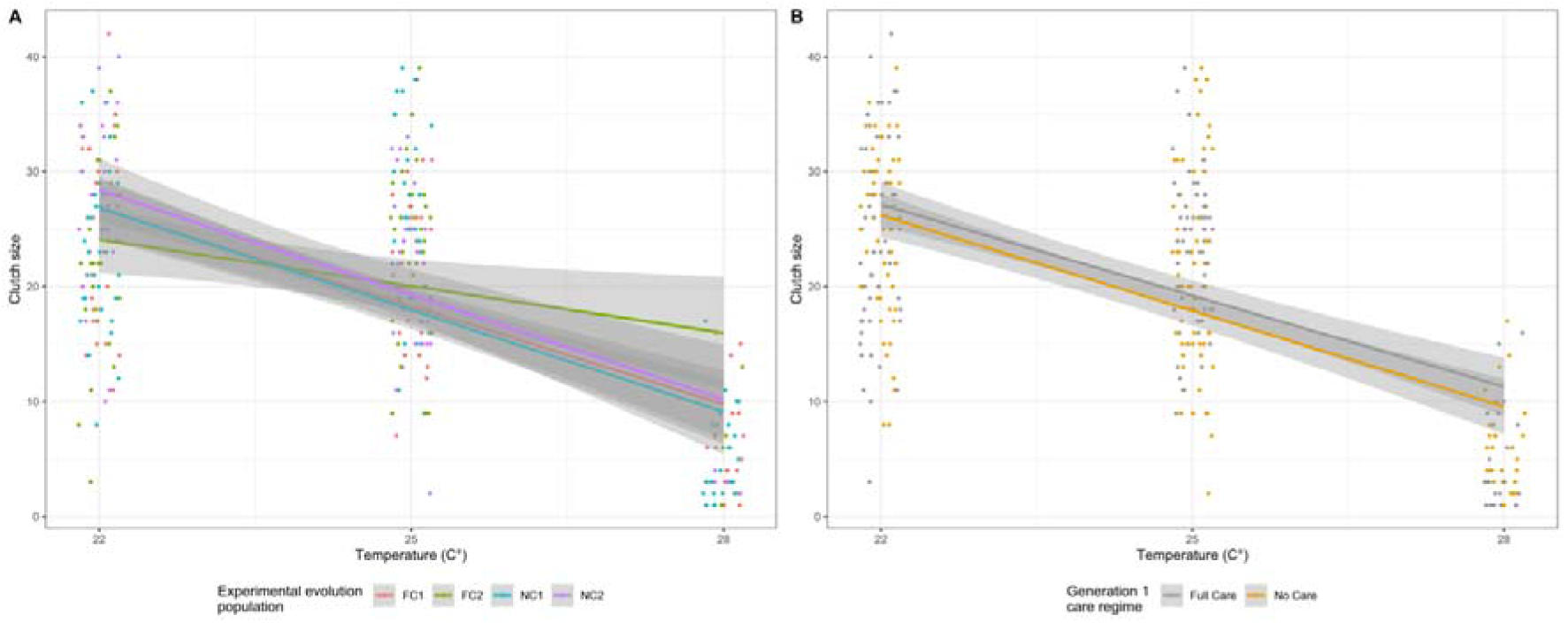
The clutch size of pairs of beetles that were kept in incubators at different levels of heat stress throughout sexual maturation in relation to A) population evolutionary history and experimental block and B) generation 1 care regime. Only pairs that laid at least one egg are presented. Points represent data for individual pairs (jittered on the x-axis), lines represent the reaction norms for each treatment and the shaded areas indicate the 95 % confidence intervals.

**Table 4.** Results of GLMs looking at variables affecting A) failure to lay: no eggs produced versus at least one egg laid and B) clutch size for pairs that laid at least one egg. Terms retained in the optimal model based on AIC_C_ are presented.

| Independent variable | Estimate | SE | z | p |
| --- | --- | --- | --- | --- |
| **A) Failure to lay** |  |  |  |  |
| Intercept | 4.488 | 0.537 | 8.360 | <0.001 |
| Block (2) | -2.082 | 0.422 | -4.933 | <0.001 |
| Temperature | -3.287 | 0.411 | -7.994 | <0.001 |

| B) Clutch size |  |  |  |  |
| --- | --- | --- | --- | --- |
| Intercept | 2.926 | 0.034 | 87.291 | <0.001 |
| Block (2) | 0.101 | 0.052 | 1.929 | 0.055 |
| Temperature | -0.350 | 0.033 | -10.568 | <0.001 |
| Block:temperature | 0.131 | 0.055 | 2.365 | 0.019 |

### Hatching failure and brood size

Neither parental population evolutionary history, nor generation 1 care regime, were retained in the optimal model of hatching success vs failure (Table S9). Parents kept at higher incubator temperatures between eclosion and sexual maturity were more likely to experience complete hatching failure (Figure 6, Tables 5A & S9) – with no eggs hatching in any of the broods where the parents had developed at 28 °C.

**Figure 6.**
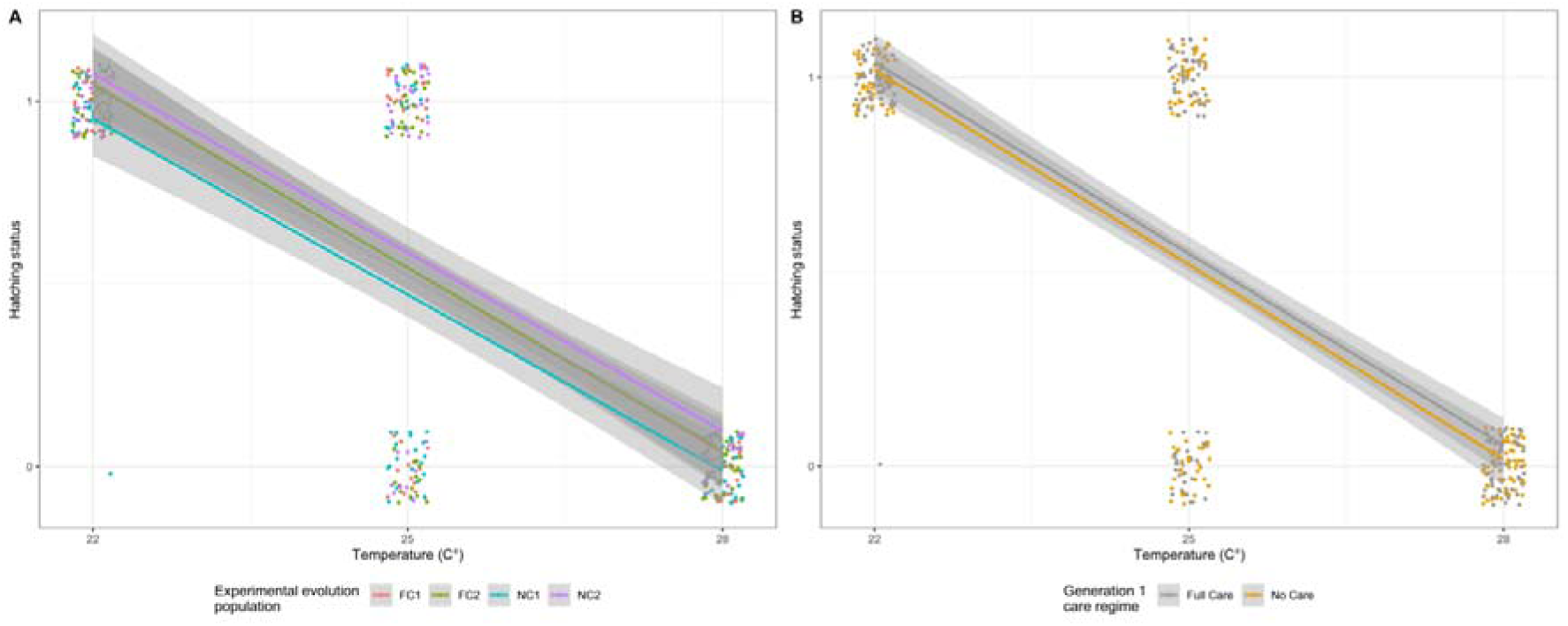
The hatching status (1 = at least 1 hatched larva, 0 = no larvae) of clutches from pairs of beetles kept in incubators at different levels of heat stress throughout sexual maturation in relation to A) population evolutionary history and experimental block and B) generation 1 care regime. Points represent data for individual pairs (jittered on the x- and y-axes), lines represent the reaction norms for each treatment and the shaded areas indicate the 95 % confidence intervals.

**Table 5.** Results of GLMs looking at variables affecting A) hatching success (at least one larva) or failure (no larvae) and B) brood size for pairs where at least one larva hatched. Terms retained in the optimal model based on AIC_C_ are presented.

| Independent variable | Estimate | SE | z | p |
| --- | --- | --- | --- | --- |
| **A) Failure to hatch** |  |  |  |  |
| Intercept | 3.091 | 0.396 | 7.817 | <0.001 |
| Temperature | -2.741 | 0.349 | -7.855 | <0.001 |

| B) Brood size |  |  |  |  |
| --- | --- | --- | --- | --- |
| Intercept | 1.878 | 0.130 | 14.405 | <0.001 |
| Block (2) | 0.289 | 0.071 | 4.093 | <0.001 |
| Temperature | -0.544 | -0.044 | -12.416 | <0.001 |
| Clutch size | 0.029 | 0.005 | 6.188 | <0.001 |

Fewer larvae hatched when their parents had been kept at 25 °C than 22 °C (Figure 7), and more larvae hatched when there were more eggs to start with (Tables 5B & S10). Additionally, for pairs where at least one larva hatched, the relationship between temperature and brood size was different between blocks, with block 1 pairs having more hatching larvae at 22 °C and block 2 pairs having more larvae at 25 °C (Figure 7B).

**Figure 7.**
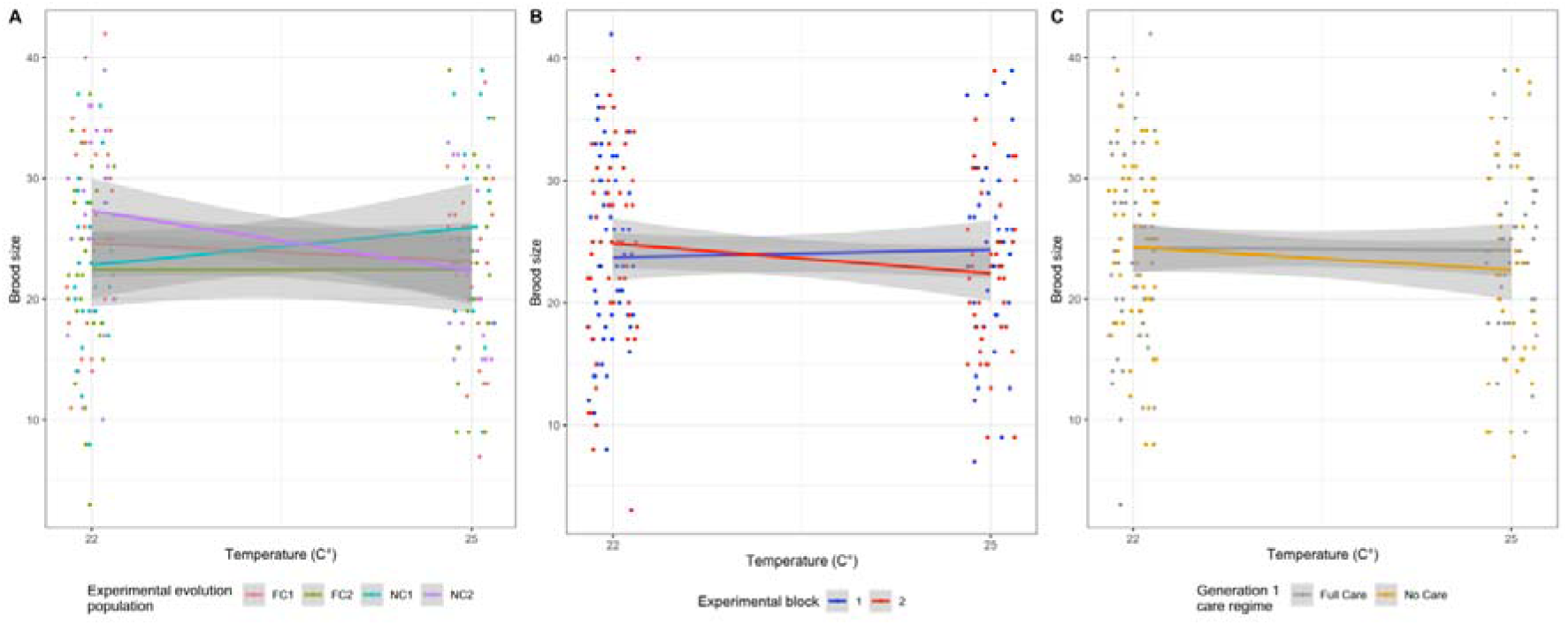
The brood size of pairs of beetles that were kept in incubators at different levels of heat stress throughout sexual maturation in relation to A) experimental evolution population, B) experimental block, and C) generation 1 care regime. Only pairs with at least one hatching larva are presented. Points represent data for individual pairs (jittered on the x- and y-axes), lines represent the reaction norms for each treatment and the shaded areas indicate the 95 % confidence intervals.

## Discussion

We tested experimentally whether a population’s past evolutionary exposure to contrasting levels of post-hatching care predicted its capacity to cope with heat stress during sexual maturation. The key findings of our experiments are summarised in Table 6, for each of the correlates of fitness we measured.

**Table 6.** The key predictors tested, their effects on correlates of fitness measured, and the hypothesis supported by the findings. “None” indicates that the predictor variable did not have a significant effect on the correlate of fitness. Full statistical results are presented in Tables 1-5.

| Predictor<br>variables<br>tested | Effect on correlates of fitness |  |  |  |  |  |  | Hypothesis<br>supported |
| --- | --- | --- | --- | --- | --- | --- | --- | --- |
|  | Survival | Lytic<br>activity | Carcass<br>sphericity | Failure to<br>lay | Clutch size | Hatching<br>failure | Brood size |  |
| Population<br>evolutionary<br>history (No<br>Care or Full<br>Care) | Lower<br>survival in<br>No Care | None | None | None | None | None | None | Beneficial<br>mutation |
| Generation 1<br>care regime<br>(No Care or<br>Full Care) | None | In males,<br>lower<br>lytic<br>activity<br>when<br>they received<br>no post-hatching<br>care as<br>larvae | Lower<br>sphericity<br>when they<br>received no<br>post-hatching care as<br>larvae | None | None | None | None | N/A |
| Temperature | Lower<br>survival at<br>higher<br>temperature<br>in block 2 | In<br>females,<br>lower<br>lytic<br>activity<br>at higher<br>temperat<br>ure | Lower<br>sphericity at<br>higher<br>temperature | Higher<br>failure at<br>higher<br>temperature | Smaller<br>clutch size at<br>higher<br>temperature,<br>particularly<br>in block 1 | Higher<br>hatching<br>failure at<br>higher<br>temperature | Smaller<br>brood size at<br>higher<br>temperature | N/A |
| Block | Lower<br>survival at<br>higher | Stronger<br>lytic<br>activity | None | Higher<br>failure in<br>block 2 | Smaller<br>clutch size at<br>higher | None | Larger<br>broods in<br>block 2 | Founder<br>effect |
|  | temperature<br>in block 2 | in block<br>2 |  |  | temperature,<br>particularly<br>in block 1 |  |  |  |

We found that populations that had evolved under Full Care experienced better survival during sexual maturation across all temperatures. This is consistent with the “beneficial mutation hypothesis”. Nevertheless the effect size was small, which could be due to exposure to relatively few generations of divergent selection and/or relatively little divergence in selection pressure since both No Care and Full Care populations developed in the relatively benign conditions of the lab.

Individuals derived from the No Care evolving populations had poor survival in the incubators, even in the most benign thermal environment, compared with those from the Full Care populations. It has previously been shown that individuals in the No Care populations carry a lower mutation load (Pascoal et al. 2022). Perhaps this means that, by chance, they lack the cryptic genetic variation that enables them to cope with new levels of thermal stress during sexual maturation (Paaby & Rockman, 2014). Alternatively, perhaps the No Care populations were intrinsically inferior by this point, and more prone to dying between eclosion and sexual maturity, regardless of the temperature to which they were exposed. The lack of a significant effect of the interaction between population evolutionary history and temperature on incubator survival is consistent with this alternative interpretation. However, in all other correlates of fitness we detected no differences between individuals drawn from the No Care versus Full Care populations, which suggests that the individuals from the No Care populations were not intrinsically inferior in each of their traits. Overall, our experimental data allow us to reject the “mutation load hypothesis” and the “selection filter hypothesis”. However, we have insufficient data to conclude we have found strong support for the “beneficial mutation hypothesis”.

Unexpectedly, we found that experimental block predicted some correlates of fitness, which could be due to a stochastic founder effect. The four distinct evolving populations were established from a single large stock population derived by interbreeding beetles from four different wild populations. The size of the breeding population was identical for each block throughout the experiment (n = 100 pairs), as was the maintenance regime, which implies that any differences between blocks must be attributable to chance differences in genetic variation when the population was founded. For example, chance differences in genetic variation could explain chance differences in the persistence of haplotypes within populations, since there were no environmental differences that could otherwise explain variation in reproductive success. In analyses of earlier generations of the experimentally evolving populations, the experimental social environment (Full Care versus No Care) explained divergence between populations to a greater extent than did block (e.g. Duarte et al., 2021; Jarrett et al., 2018a, 2018b; Schrader et al., 2015c, 2015a, 2015b, 2017, 2017). By generation 46, however, the situation had reversed: block outweighed experimental social environment in explaining the response to heat stress exposure during sexual maturation.

Founder effects have long been suggested to explain the success of translocation and captive breeding in conservation programmes (e.g. Brown et al., 2009; Dolman et al., 2021; Miller et al., 1999). However, in analyses of wild populations it is often hard to disentangle the scale of founder effects from confounding factors of environmental differences (however subtle) in the new location (Kolbe et al., 2012; Szűcs et al., 2017) especially when it is difficult to monitor populations systematically after their release into the wild (Armstrong & Seddon, 2008; Mock et al., 2004; Seddon et al., 2007; Weeks et al., 2011). Nevertheless, there is some evidence that founder effects influence the response to selection in wild populations. When *Anolis sagrei* individuals with long hindlimbs were translocated to seven Bahamian islands with a vegetation structure known to favour short hindlimbed individuals, they evolved shorter hindlimbs over four years. However the extent of morphological change was attributable to founder effects (Kolbe et al., 2012). Perhaps a similar pattern explains the results we have found in this study. More broadly our study indicates the importance of the initial population size in captive breeding programmes. It also reinforces the wisdom of using genetic screening to determine whether the founding population is representative of the source population and has high heterozygosity (Bragg et al., 2021; Scott et al., 2020), especially in cases where there will be no migration into the population, such as on an isolated island. Our findings are of particular interest from a practical perspective considering that many conservation captive breeding programmes have, by necessity, used smaller founder populations than ours (Witzenberger & Hochkirch, 2011).

However, for some of the traits we measured (failure to lay, clutch size, failure to hatch) we found that temperature had such a strong negative effect that it outweighed any effects that could be attributed to the population’s genetic background. This is consistent with previous work across different species of burying beetle, which found that high temperatures during breeding led to reduced clutch and brood sizes and a greater likelihood of complete brood failure (Keller et al., 2021; Moss & Moore, 2021; Ney, 2021; Ong, 2019; Quinby et al., 2020; Sun & Kilner, 2020). In all of these studies the extreme temperature treatment continued throughout breeding, whereas in this study we elevated temperature only during sexual maturation to better simulate the natural history of the burying beetle. Our results suggest that there are sensitive windows in development which affect breeding success even when reproduction occurs under benign conditions. In a world with an increasing frequency and intensity of heat waves (Perkins-Kirkpatrick & Lewis, 2020), this study shows that an extreme temperature event can reduce breeding success even if it is only brief, as long as it coincides with these sensitive windows in development.

There are two likely causes for a reduction in egg laying and egg hatching at higher temperatures: 1) behavioural – beetles maturing at higher temperatures mated less, and/or 2) physiological – beetles maturing at higher temperatures had underdeveloped gonads. Particularly in males, exposure to high temperatures prior to pairing leads to reduced courtship behaviour in *Aphidius avenae* (Roux et al., 2010) and *Drosophila melanogaster, Drosophila simulans and Drosophila mojavensis* (Patton & Krebs, 2001). There is also a great deal of evidence to suggest that thermal stress during sexual maturation can have a physiological effect on insects. In *Drosophila melanogaster* high temperatures during development cause a range of spermatid cytological abnormalities (David et al., 2005; Rohmer et al., 2004) and in *Tribolium castaneum* cause lower testis volume (Sales et al., 2021). In females, higher temperatures during development cause smaller oocytes in *Pararge aegeria* (Gibbs et al., 2010) and cause *Bicyclus anynana* to lay more, smaller eggs (Steigenga & Fischer, 2007). From this study we are unable to say whether the effects were behavioural, physiological or both, and whether it was the males or females that drove the patterns of reduced reproductive fitness at higher temperatures. Further investigations tracking copulatory behaviour, mating thermally stressed males with control females and vice versa, and dissecting ovaries and testes following heat stress are needed to answer these questions.

It is widely accepted that a population’s standing genetic variation will predict its response to new environmental challenges. Our study suggests it is possible that the evolutionary history of burying beetle populations might predict the response to heat stress during sexual maturation (because No Care populations had lower survival at all temperatures in the incubators), although further work is needed before a robust conclusion can be drawn. However, stochastic genetic variation, due to founder effects or genetic drift, was potentially a more powerful predictor of population resilience upon exposure to new thermal environments. Translocation and captive breeding programmes should be designed accordingly to maximise their chance of success.

## Supporting information

Cover page

