## Supplementary material for "Can recent evolutionary history promote resilience to environmental change?": Cover page

**Running title:** Evolutionary history in resilience to change

**Author names and affiliations:**

Eleanor K. Bladon^1^*, Sonia Pascoal^1,2^ & Rebecca M. Kilner^1^

^1^Department of Zoology, University of Cambridge, Downing Street, Cambridge, CB2 3EJ, UK.

^2^Department of Haematology, University of Cambridge, Long Road, Cambridge, CB2 0PT, UK.

**Author contributions:** EKB, RMK and SP designed the experiments; EKB and SP collected the data; EKB analysed the data; EKB and RMK wrote and edited the manuscript.

**Funding:** This project was supported by a Consolidator’s Grant from the European Research Council (310785 Baldwinian_Beetles), a Wolfson Merit Award from the Royal Society, The Leverhulme Trust (RPG-2018-232) and The Isaac Newton Trust (18.23(q)), each to RMK. EKB was funded by a Biotechnology and Biological Sciences Research Council PhD studentship (BB/M011194/1).

**Acknowledgements:** We thank Chris Swannack and Sue Aspinall for helping with beetle maintenance.

**Conflict of interest:** The authors declare no conflicts of interest

**Data archiving:** Data will be made available on Dryad at the time of publishing
